# Inhibiting the interaction between the mitochondrial receptor Tom70 and SARS CoV 2 Orf9b with small molecules

**DOI:** 10.64898/2026.04.27.721040

**Authors:** CJ San Felipe, Kliment A Verba, Nevan J Krogan, Michael Grabe, James S Fraser

**Affiliations:** Department of Bioengineering and Therapeutic Sciences, University of California, San Francisco, San Francisco, United States; Department of Cellular and Molecular Pharmacology, University of California, San Francisco, San Francisco, United States; Quantitative Biosciences Institute, University of California, San Francisco, San Francisco, United States; Department of Pharmaceutical Chemistry and Cardiovascular Research Institute, University of California, San Francisco, San Francisco, United States; Gladstone Institute of Data Science and Biotechnology and Gladstone Infectious Disease Institute, J. David Gladstone Institutes, San Francisco, CA 94158, USA; Quantitative Biosciences Institute (QBI) COVID-19 Research Group (QCRG), San Francisco, CA 94158, USA

## Abstract

The SARS CoV 2 accessory protein Orf9b is in a complex monomer-dimer equilibrium that influences its interactions with the host mitochondrial receptor Tom70. This interaction is critical for viral suppression of a Type-1 interferon response during infection. Modulating this equilibrium with a small molecule, either by stabilizing the Orf9b homodimer or blocking its interaction with Tom70, represents a promising strategy for restoring interferon signaling and the antiviral response. To build tool molecules that could test this concept, we performed two screens: a crystallographic fragment screen against the Orf9b homodimer and a high-throughput fluorescence polarization screen for competitors of an Orf9b-derived peptide binding to Tom70. Fragment screening revealed two binding sites with potential to be developed into an inhibitor: one located at the peripheral dimer interface and the other just outside the lipid-binding channel that defines the central dimer interface. Functionalization of the fragments outside of the lipid-binding channel with hydrophobic moieties stabilized the Orf9b homodimer thereby indirectly inhibiting association with Tom70. In parallel, the high throughput screen for competitive inhibitors of the Tom70:Orf9b interaction discovered a separate series of molecules. These molecules display dynamic structure activity relationship (SAR) and could be improved in the future to modulate the interaction between Tom70 and potentially a wide range of substrates. Collectively, these results demonstrate the feasibility of two distinct strategies to manipulate the Orf9b-Tom70 equilibrium, which is critical to the host response to SARS CoV 2 infection.

## Introduction

During viral infection, one of the necessary steps to successful replication is the suppression and evasion of the host’s immune system (Stetson and Medzhitov 2006; Takeuchi and Akira 2009). The virus SARS CoV 2 accomplishes this in several ways by encoding accessory viral proteins that help to antagonize immune responses (Wang et al. 2021; Han et al. 2022, 2021; Shemesh et al. 2021; Felgenhauer et al. 2020). One of those factors is Orf9b. Orf9b (Open Reading Frame 9b) is encoded by both SARS CoV and SARS CoV 2 through an alternative open reading frame within the Nucleocapsid (N) gene (Xu et al. 2009; Yang et al. 2023; Meier et al. 2006).

Studies of Orf9b’s role in viral infection going back to SARS CoV showed that Or9b localized to the mitochondrial membrane (Meier et al. 2006). Later work investigating SARS CoV 2 revealed that Orf9b could suppress a Type 1 interferon response (Jiang et al. 2020). It was not until the emergence of SARS CoV 2 and the subsequent Covid 19 pandemic that structures of Orf9b bound to host factors emerged providing insights into the relationship between Orf9b’s structure and its role in viral infection: Cryo-EM (Gordon et al. 2020) and X-ray structures (Gao et al. 2021) revealed that Orf9b bound to the host factor Translocase of Outer Membrane 70 (Tom70), a mitochondrial outer membrane receptor involved in the import and biogenesis of cytoplasmically translated protein to the mitochondria (Backes et al. 2021, 2018).

Tom70 fulfills two important functions: it facilitates the import and biogenesis of cytoplasmically translated proteins to the mitochondria through its chaperone binding activity and matrix targeting sequence (MTS) recognition and it also supports the activation of the type-1 interferon response through its association with several signaling proteins including MAVS and TBK1 (Lin et al. 2010; Wei et al. 2015; Liu et al. 2010; Hansen and Herrmann 2019; Diekert et al. 1999). The soluble domain of Tom70 possesses a C-terminal binding site that is involved in binding to targeting sequences of cytoplasmically translated proteins that fold into amphipathic alpha helices for import into the mitochondria (Backes et al. 2021). Both structures of the Orf9b:Tom70 complex showed that Orf9b can adopt a monomeric amphipathic helical fold when bound to Tom70 at the C-terminal binding site and to date are the only structures of Tom70 with a substrate bound to the C-terminal site (Young et al. 2003; Backes et al. 2018; Gordon et al. 2020).

In support of fulfilling two different functions (mitochondrial import and innate immune signaling), Tom70 also possesses a second binding site near its cytoplasmically exposed N-terminus for binding chaperones like Hsp70/90 (Backes et al. 2018; Young et al. 2003). In the context of mitochondrial import, both sites have been shown to be important, likely because the substrate proteins that are imported by Tom70 are membrane proteins that have exposed hydrophobic regions which require chaperones to remain soluble (Backes et al. 2021). In the context of innate immune signaling, the role of the C-terminal binding site is unclear, however, point mutations of the N-terminal chaperone binding site have been shown to block Tom70’s association with chaperones such as Hsp90 which is sufficient to inhibit innate immune signaling (Liu et al. 2010).

While several studies have suggested how Orf9b could antagonize innate immune signaling through Tom70, the exact mechanistic details remain unclear. Thorne et al demonstrated that Orf9b binding to Tom70 can be inhibited through phosphorylation of Orf9b by a host kinase that blocks binding to the C-terminal binding site on Tom70 and restores interferon signaling (Thorne et al. 2022) suggesting a role for the C-terminal binding site in innate immune signaling. Structural work by several groups have reported that binding of Orf9b at the C-terminal binding site results in slight rearrangements of critical residues in the N-terminal binding site that are responsible for binding to the EEVD motif on chaperones (Gordon et al. 2020; Gao et al. 2021; Bachochin et al. 2026). Bachochin et al were the first to solve the structure of apo-hTom70 by crystallography which demonstrated that Orf9b induced a conformational change in the C-terminal domain of Tom70 upon binding. Prior investigation by Gao et al using peptides derived from Orf9b and Hsp90 suggested that when bound at the C-terminal binding site on Tom70, Orf9b allosterically modulates the N-terminal binding sites affinity for peptides derived from chaperones as measured by ITC. Conversely, a previous study using yeast homologs of Tom70 had shown that chaperone-derived peptides do not affect the binding of proteins to the C-terminal binding site (Mills et al. 2009) suggesting a unidirectional allosteric relationship between C and N-terminal domains. The allosteric hypothesis has been challenged by a recent paper that has proposed that the primary driver of chaperone binding inhibition is not due to an allosteric mechanism but due to Orf9b competitively blocking chaperone binding while associated with Tom70 at the C-terminal binding site (Sherer et al. 2025). In this paper, the authors show that the intrinsically disordered C-terminus of Orf9b (which is not resolved in either structure of the Orf9b:Tom70 complex) can extend down from Tom70’s C-terminal binding site (where Orf9b adopts a helical conformation) and obstruct Tom70’s N-terminal binding site to inhibit chaperone binding and thus interferon signaling (Sherer et al. 2025)

We have previously demonstrated that Orf9b exists in a homodimer form that is in a coupled binding equilibrium with a 1:1 Tom70:Orf9b complex (San Felipe et al. 2025). X-ray structures of both SARS CoV and CoV-2 show Orf9b homodimers adopt a beta sheet rich confirmation with a central channel that is occupied by a lipid-like ligand (Meier et al. 2006; San Felipe et al. 2025). The Orf9b homodimers must dissociate into alpha-helical Orf9b monomers that go on to bind Tom70 leading to the suppression of a type-1 interferon response (**Figure 1A**) (San Felipe et al. 2025). We also demonstrated that this equilibrium is regulated by lipid-binding to the Orf9b homodimer which stabilizes the homodimer conformation approximately 100-fold relative to the apo-homodimer and substantially slows the equilibrium with Tom70 (San Felipe et al. 2025), which demonstrates a second possible source of regulation for the Orf9b-Tom70 interaction.

**Figure 1:**
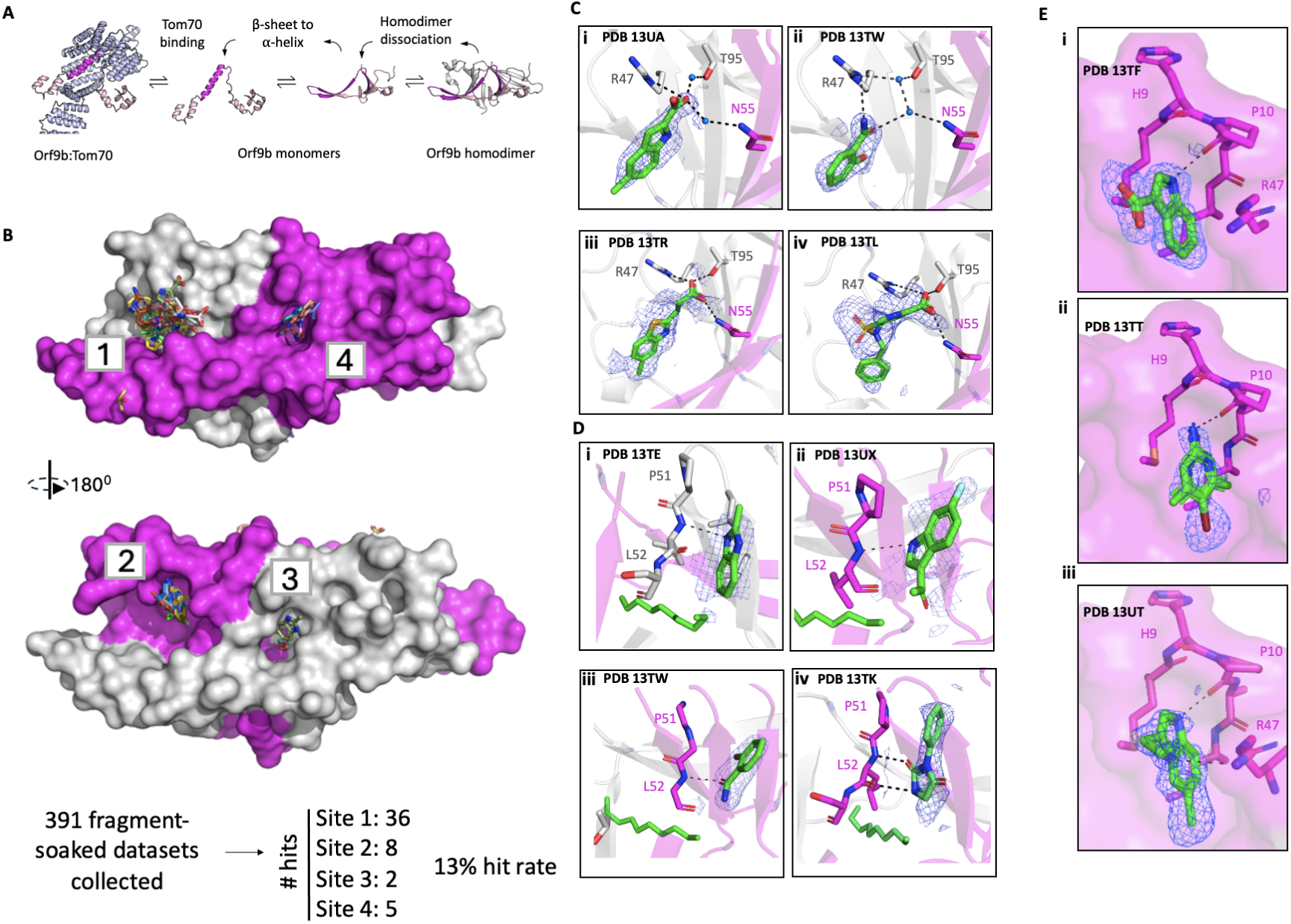
Overview of Orf9b homodimer crystallization system and fragment screening. (A) Model for the Tom70-Orf9b equilibrium. Orf9b homodimer dissociates into Orf9b monomers which undergo a conformational change from *β*-sheet to *α*-helix. *α*-helical Orf9b binds to Tom70. (B) Overview of Orf9b homodimer with modeled fragment hits. Orf9b homodimer surface is modeled with fragments as sticks including the number of modeled hits per binding site. Hydrogen bonds are shown as black dashes with a 4Å cutoff. (C) Overview of binding site 1 showing residue side chains. Representative binding poses of modeled fragments bound at site 1. PanDDA event maps are shown contoured to 2*σ*. Hydrogen bonds are shown as black dashes with a 3.5Å cutoff. (D) Representative binding poses of fragments identified at sites 3 and 4 corresponding to the homodimer central channel, the lipid molecule colored in green is modeled to provide orientation. PanDDA event maps are shown contoured to 3*σ*. Hydrogen bonds are shown as black dashes with a 3.5Å cutoff. (E) Overview of binding site 2 showing representative fragments with PanDDA event maps contoured at 2*σ*. Fragment in structure PDB 13TT (ii) is shown with two conformations.

To date, there are no known chemical tools or therapeutics targeting either Orf9b (in either its monomeric or homodimeric form) or Tom70. Small molecules targeting either Orf9b or Tom70 could potentially be used to probe the role of the Orf9b homodimer as well as the mechanism of Tom70-mediated interferon suppression. Several published reports on SARS CoV 2 viral factors have also highlighted how developing therapeutics against these proteins can help illuminate the mechanisms that underlie their function and role in viral pathogenesis as well as the host processes they antagonize (Suryawanshi et al. 2025; Detomasi et al. 2025; Van Damme et al. 2025; Meyer et al. 2025). In this work, we performed two distinct screens to develop chemical matter targeting either Orf9b homodimers and Tom70 to better explore the Orf9b:Tom70 interaction. First, we used crystallographic fragment screening to explore the targetable space of the Orf9b homodimer and developed PEG-functionalized small molecules that “glue” the homodimer together to kinetically slow its association with Tom70. Next, we used a high-throughput fluorescence polarization peptide displacement assay to identify Tom70-binding small molecules that compete for Orf9b binding at the C-terminal binding site. Collectively, these results represent two promising strategies for restoring the host immune response by modulating the Orf9b:Tom70 association by targeting either the viral fold-switching process or the direct interaction of a viral protein with the host machinery.

## Results

### Developing an Orf9b homodimer crystallization system for fragment screening

We sought to identify crystallization conditions that would both yield reproducible crystals that diffracted to high resolution and be amenable to DMSO for soaking fragment libraries (Collins et al. 2018). We optimized crystallization conditions for the Orf9b homodimer based on an existing structure of Orf9b from Protein Data Bank (PDB 6Z4U) which yielded crystals that diffracted to an average resolution of 1.8Å with minimal changes to unit cell dimensions (**Figure 1–figure supplement 1A**). Orf9b crystallizes in a P212121 space group with two copies of Orf9b in the asymmetric unit forming the homodimer conformation. Orf9b purifies as a homodimer with a co-purifying lipid bound in the central hydrophobic channel (San Felipe et al. 2025). Although we have shown that it is possible to refold the Orf9b homodimer to eliminate the lipid from the central channel, we found that apo-Orf9b homodimer crystals were far less tolerant to DMSO and on average diffracted to resolutions worse than 2.5Å. Importantly, we found that microseeding helped increase crystal reproducibility and yielded larger crystals on average. Therefore, for our crystallographic fragment screening campaign we elected to use the lipid-bound protein to identify additional potential binding sites that could be exploited to stabilize the Orf9b homodimer.

We conducted DMSO tolerance tests on Orf9b homodimer crystals and found that these crystals were highly tolerant up to 20% DMSO for 24 hours. One of the principal advantages of screening fragments is that they can more efficiently sample chemical space and bind to regions of a protein that larger lead-like compounds cannot access, however, their small size also tends to result in compounds with very low affinity (mM to high uM) (Murray and Rees 2009). Coupled with the nature of soaking compounds into crystals, fragments may not bind at high occupancy which results in weak electron density in conventional electron density maps to support their placement in the structure. To resolve fragments weakly bound to the Orf9b homodimer, we utilized the Pan-Dataset Density Analysis (PanDDA) method (Pearce et al. 2017). Before we began our fragment soaking screen, we first generated a set of DMSO-soaked structures which we would use for calculating the background maps which diffracted to between 1.5-1.8Å and were isomorphous (**Figure 1–figure supplement 1B**). We next soaked two fragment libraries into Orf9b homodimer crystals: the 320 fragment Enamine essential library and a 91 compound in-house library (UCSF_91) (Schuller et al. 2021).

### Fragments bound in 4 hotspots, representing 2 biologically equivalent binding sites

After soaking, we collected 391 fragment-soaked datasets which diffracted to between 1.26Å and 2.46Å (average 1.8Å) resolution and observed that soaking fragments in DMSO into our crystals did not alter the unit cell properties (**Figure 1–figure supplement 1C**). We used PanDDA to generate event maps that allowed us to model 51 fragments bound to the Orf9b homodimer (**Figure 1B**), representing a hit rate of approximately 13%. These fragments were spread out over 4 binding sites which were defined as binding sites 1, 2, 3, and 4 (**Figure 1B**). As Orf9b homodimer is a symmetric protein, binding site 1 is biologically identical, but crystallographically distinct, to site 2; the same is true for sites 3 and 4. Sites 1 and 2 sit at a peripheral dimer interface whereas sites 3 and 4 sit at the central lipid-binding channel entrance.

We identified 36 fragments bound to site 1, with an additional 8 fragments bound at the biologically equivalent site 2, which we refer to collectively as the “peripheral dimer interface” (**Figure 1C and E**). The peripheral dimer interface is largely composed of nonpolar and hydrophobic residues with a patch of polar residues composed of R47 and T95 on chain A and N55 and S53 on chain B (**Figure 1C**). Most fragments at site 1 leverage hydrogen bonds with either R47 on chain A or N55 on chain B as well as water molecules to bridge hydrogen bonds with chain B of the Orf9b homodimer (**Figure 1Ci and Cii**).

Despite sites 1 and 2 being biologically identical, we observed that there were differences in the local crystal environment. First, we observed that both binding sites are composed of the two chains of Orf9b but also have a neighboring symmetry mate that encloses a bipartite pocket (**Figure 1–figure supplement 2A**). While the relative positions of the symmetry mates with respect to sites 1 and 2 are largely the same, we observed that site 1 is slightly more open than site 2. At site 2, the neighboring symmetry mate is packed more tightly against the binding site, reducing its size (**Figure 1–figure supplement 2A**). We also observed additional electron density in site 2 that was present in both fragment bound and DMSO soaked datasets which we ascribed to a molecule of DMSO (**Figure 1–figure supplement 2E**). This added density is absent in site 1 and further reduces the size of binding site 2 while also constraining the fragments binding orientation (**Figure 1–figure supplement 2E**). In binding site 2, we observed that R47 of chain B can be displaced by fragments (**Figure 1–figure supplement 2D**). This conformational change was not observed in the biologically identical site 1. The R47 conformational change in site 2 opens up access to the backbone between residues P10 and A11 which allows for hydrogen bonding between the fragment and the amide carbonyl on P10 and the backbone nitrogen between A11 (**Figure 1–figure supplement 2D**). This results in fragments at site 2 binding in a different orientation compared to site 1 (**Figure 1–figure supplement 2B**). Therefore, although site 1 contains fragments that make contacts that are exclusive to the two chains of Orf9b in the homodimer, site 2 may be producing artifactual binding poses that are dictated by the neighboring symmetry mates. We therefore did not further pursue optimization strategies to improve fragment binding at site 2 and assumed that site 1 is more representative of how compounds would bind to the homodimeric assembly of Orf9b in solution.

We next turned our attention to the fragment binding events near the homodimer central channel. As previously mentioned, our crystallization system for the Orf9b homodimer contains a co-purifying lipid that is bound in the central channel formed by the Orf9b homodimer, and we did not observe any fragments that displaced the lipid. We did observe several fragments that were bound near either end of the central channel entrance with the majority of fragments bound near the entrance formed by chain B which we termed site 4 and 2 fragments bound at the opposite (but biologically identical) entrance which we termed site 3 (**Figure 1D**). In all instances, fragments at site 3 and 4 make either one or 2 hydrogen bonds with the backbone of either chain A or B. As with sites 1 and 2, we also inspected the crystal environment around sites 3 and 4 to determine if neighboring crystal contacts influences the distribution of hits and their binding poses (**Figure 1–figure supplement 2C).** We found that for site 3, the nearest symmetry mate was approximately 18Å away from the channel entrance whereas site 4 has a symmetry mate that packs close to the opposite end of the channel (**Figure 1–figure supplement 2C**).

### Conservative fragment growing of external Orf9b homodimer interface yields weakly binding analogs

We sought to improve upon our original fragment hits with the goal of stabilizing Orf9b in the homodimer conformation to inhibit binding to Tom70. Given that most of our fragments were observed binding to site 1, we first considered improving compounds that targeted this binding site to determine the suitability of developing lead-like compounds that could stabilize the homodimer. We pursued a conservative fragment growing strategy given the size of the binding site and that most of our hits overlapped with each other. In parallel to our fragment hits, we turned to FTMap to explore the binding site and determine what types of growing strategies could be tolerated (Brenke et al. 2009). FTMap consensus sites confirmed that sites 1 and 2 where we observed fragments bound on the homodimer surface are predicted hot spots for fragment binding with FTMap consensus site 1 chemical probes following a similar binding orientation as our crystal structures (**Figure 2–figure supplement 1A**). While FTMap consensus site 2 was also predicted to be a hot spot like our observed fragments bound at site 2, the binding orientation was noticeably different than site 1 but also varied from the experimentally observed fragment binding poses in our crystallographic data sets (**Figure 2–figure supplement 1B**). We superimposed one of our representative site 1 fragment-bound structures to the FTMap consensus site (PDB 13UB) (which coincided with site 1) and treated the consensus probe clusters as mapping out the accessible area of site 1 for conservative fragment growing.

**Figure 2:**
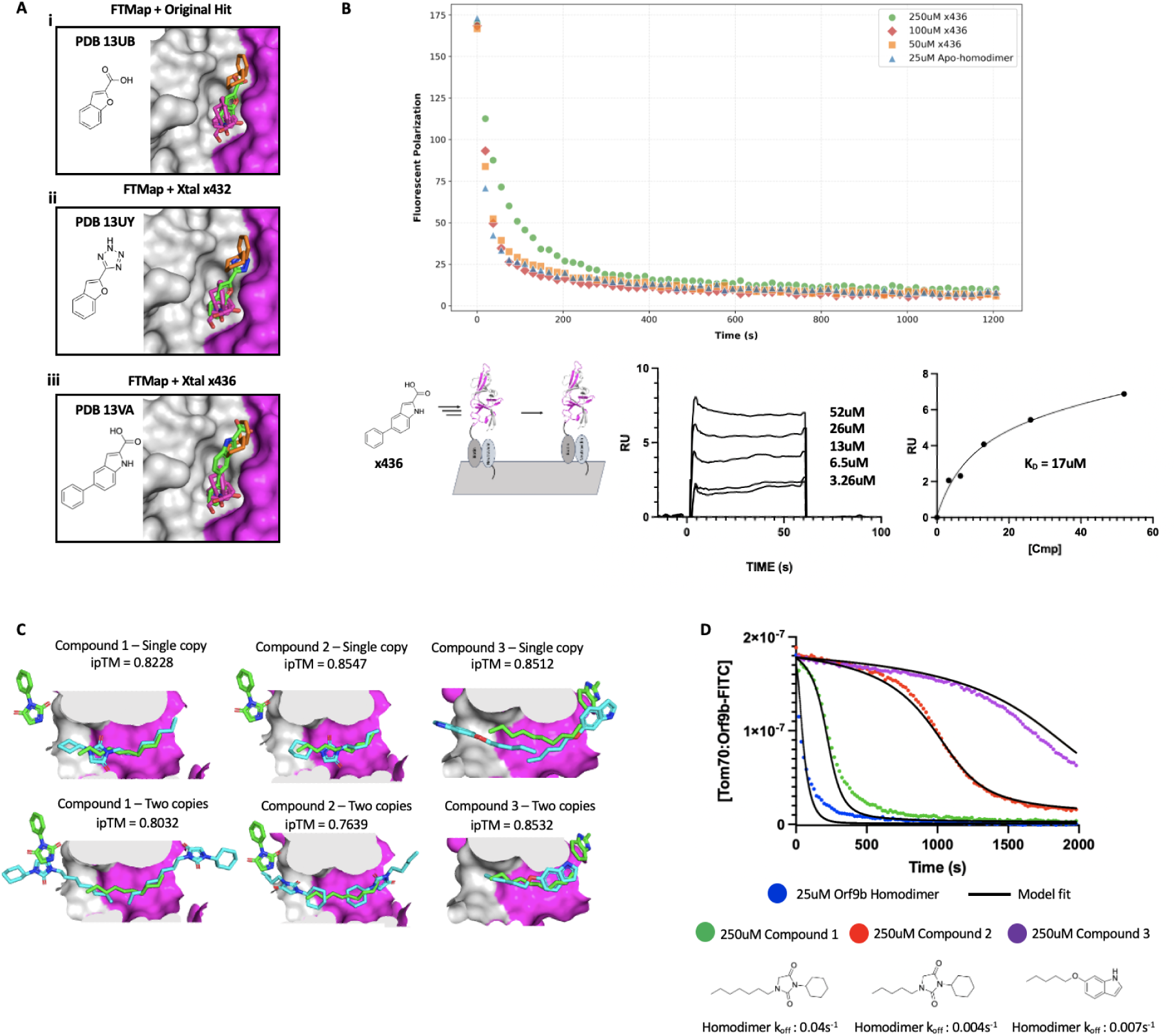
Overview of Orf9b homodimer analog binding and binding behaviors. (A) Structurally resolved fragment hit and analog compounds bound at site 1 with FTMap chemical probes superimposed. PDB codes for deposited structures are displayed in the top left corner of each panel. (B) (Top) Kinetic traces of Orf9b homodimer binding to Tom70 in the presence of increasing x436 analog concentration. (Bottom left) Diagram of SPR experiment with immobilized Orf9b homodimer and x436 analog compound to measure binding at external dimer interface. (Bottom middle) Surface plasmon resonance sensorgram showing x436 analog compound binding to the Orf9b homodimer immobilized on the surface. (Bottom Right) Modeling SPR sensorgram equilibrium response with a non-linear regression to determine the affinity for the Orf9b homodimer of 17uM. (C) (Top) Chai1 predictions of Orf9b homodimer co-folded with one copy of lipidated analog and ipTM scores in the Orf9b homodimer central channel. (Bottom) Chai1 predictions of Orf9b homodimer co-folded with two copies of the lipidated analog and ipTM scores. The original fragment and modeled lipid are highlighted in green with the Chai1 predicted analogs in cyan. (D) Kinetic model overlay to FP competition kinetic assay using Orf9b homodimer incubated with lipidated analog as the competitor. Structures of lipidated analogs that exhibit slow homodimer dissociation.

We tested strategies for expanding upon our initial fragment hit (PDB 13UB) by either growing from the aromatic ring towards the non-polar residue patch, or, extending off of the carboxylic acid group. When superimposed with the FTMap probe clusters, we reasoned that aromatic groups similar in size to benzene could be tolerated. We also hypothesized that this could displace water molecules that form bridging hydrogen bonds between the fragments and the patch of polar residues to improve affinity. We incorporated these considerations into the design of analog compounds with the goal of improving the specific bonds formed between the fragments/analogs and the two chains of Orf9b (**Figure 2A**). We purchased the closest analog compounds based on these designs and soaked them into crystals. Just as with the parent fragment, the analog compounds were observed binding in site 1 as well as in the same orientation as the original hit, confirming that the conservative changes we made retained binding to the Orf9b homodimer (**Figure 2A**).

We next sought to determine the affinity of our analogs for the homodimer using SPR. We immobilized the biotinylated (lipid-bound) homodimer and tested both the original fragment hit and our analog compounds that we structurally resolved bound to the Orf9b homodimer (**Figure 2B**). Unsurprisingly, the original fragment hit exhibited no detectable binding to the Orf9b homodimer, which is characteristic for fragments with very low affinities (Keserű et al. 2016). We next tested analogs that we structurally resolved bound to the homodimer to determine if conservative growing strategies yielded any improvement in affinity by SPR. We were able to obtain binding of the x436 analog to the Orf9b homodimer which exhibited fast kon and koff which made kinetic fitting unsuitable. We instead plotted the average equilibrium response against the compound’s concentration to determine the Kd. While we were not able to saturate the response, due to the solubility limits of the compound, we can fit an estimated Kd of 17uM for the x436 analog to the Orf9b homodimer (**Figure 2B**). Based on our SPR data, we next investigated if the x436 analog could stabilize the Orf9b homodimer through the external dimer interface in an FP competition kinetic binding assay that we have previously described (San Felipe et al. 2025). We tested three concentrations of the compound from 50uM to 250uM but observed no noticeable increase in homodimer stability as indicated in a shift in the kinetic binding curves compared to the apo-homodimer (**Figure 2B**). This suggests that although these compounds can bind to the external dimer interface, it may not be suitable for exercising a stabilizing effect on the Orf9b homodimer conformation. Taken together, these results suggest that although these sites are hotspots for fragment binding, the opportunities for improving upon these hits to achieve tighter affinities (and greater stabilization of the homodimer) may be very limited. Therefore, we next turned our attention to the lipid binding channel.

### Lipidated analogs exhibit increased Orf9b homodimer stability in an FP kinetic assay

We previously showed that lipid binding to the Orf9b homodimer can tightly stabilize the homodimer conformation and slow the rate of dissociation into monomers that bind to Tom70 by ∼100 fold (San Felipe et al. 2025). Presumably, the binding of lipids in this channel (which is composed of the two Orf9b chains) acts like a molecular glue to keep the homodimer conformation stabilized. Inspired by our previous work, we were interested in seeing if it was possible to develop the fragments that we observed binding near the central channel entrance into compounds that recapitulate the lipid-bound homodimer behavior. Starting from the fragments that we observed binding near the homodimer channel entrance at sites 3 and 4, we could observe that they were in close proximity to the lipid bound in the central channel of the homodimer. This presented an opportunity to merge the fragments with the lipid tail to improve both affinity as well as stability of the homodimer when bound. Inspection of our FTMap predictions further motivated us to prioritize lipid-like moieties as the FTMap model did not identify any fragment clusters larger than amides and ethanes, likely due to the physical constraints of the channels diameter (**Figure 2–figure supplement 1D**). In previous Orf9b homodimer structures, the central channel lipid was modeled as an 8-carbon alkane based on agreement with the electron density (San Felipe et al. 2025, Meier et al. 2006). We purchased several analogs based on the parent fragment with variable tail lengths that we hypothesized would point into the central channel of the apo-homodimer and increase affinity largely through hydrophobic interactions but with the added specificity that would arise from the fragment derived portion of the analogs.

We utilized our FP kinetic assay to determine if lipidated-analog binding to the Orf9b homodimer had any noticeable effects on the stability of the Orf9b homodimer and the subsequent Orf9b-Tom70 equilibrium. We first considered whether the lipidated-analogs had any binding activity for Tom70 on its own and whether changes in FP signal were due to binding to the Orf9b homodimer. We tested the compounds on their own in the presence of the pre-equilibrated Tom70:Orf9b-FITC complex and observed no significant decrease in FP signal indicating that the lipidated-analogs do not directly bind to Tom70 to drive a decrease in FP signal (**Figure 2–figure supplement 2A**). We next tested whether these compounds acted on the Orf9b homodimer specifically or if they could inhibit the Orf9b monomer from binding to Tom70 by incubating compounds with an Orf9b peptide that acts as an obligate monomer. When chasing with the pre-equilibrated Tom70:Orf9b-FITC complex, we observed no significant differences in kinetics of Orf9b peptide binding compared to the peptide on its own which showed a fast single-exponential kinetic decay in signal suggesting that these compounds do not act on the Orf9b monomer to inhibit binding to Tom70 (**Figure 2–figure supplement 2B**). Next, we took the apo-homodimer that we isolated by refolding the recombinantly expressed Orf9b (see methods) and incubated our lipidated-analogs to allow for binding in the central hydrophobic channel formed by the homodimer. We chased the pre-equilibrated Orf9b homodimer:lipidated-analog solution with the pre-equilibrated Tom70:Orf9b-FITC fluorescent complex to monitor the decrease in fluorescent polarization signal upon displacement of the Orf9b-FITC probe.

We found three lipidated-analogs out of 12 tested that exhibited a slow initial decrease in signal compared to the apo-homodimer condition which is indicative of a shift in the Orf9b-Tom70 equilibrium towards the homodimer conformation (**Figure 2D**). The failure of 9 compounds to slow the equilibrium suggests that this is not a general effect of Orf9b binding to an aliphatic tail, but rather the combined action of the specific fragment and tail portion together. Importantly, across all conditions the concentration of the Orf9b homodimer was kept constant, therefore, we hypothesized that changes in the FP kinetics observed must be driven by changes to the overall stability of the homodimer driven by lipidated-analog binding. Compared to the experimentally determined homodimer dissociation rate, we modeled the lipidated-analog conditions using a kinetic model based on ordinary differential equations that fully describe the coupled binding equilibrium between Orf9b homodimers, monomers, and Tom70 that we previously described (San Felipe et al. 2025). We showed with this model that shifts in the measured decrease in fluorescent polarization signal over time can be explained by decreases in the rate of Orf9b homodimer dissociation into monomers upon lipid binding to the homodimer conformation. We adapted this model to also determine the effect our lipidated-analogs had on the dissociation rate of the Orf9b homodimer. We modeled our lipidated-analog FP data by varying the rate of homodimer dissociation that produced the best agreement with the measured data while fixing all other rates in the equilibrium. This resulted in the rate of homodimer dissociation varying between 8.75 and 50-fold slower than the apo-homodimer when lipidated analogs are pre-incubated with the homodimer (**Figure 2D**). Compound 3 which exhibited the greatest shift in the kinetic binding curve when incubated at 250uM results in a modeled homodimer dissociation rate ∼50 times slower than the apo-homodimer. While this rate is still 7 fold faster than the lipid-bound homodimer, these results suggest that the effect exercised by these lipidated-analogs is specific to the Orf9b homodimer and that these analogs can stabilize the Orf9b homodimer to slow formation of the Orf9b:Tom70 complex.

### Chai-1 predicts lipidated-analogs bind in Orf9b homodimer central channel with varying poses

The three lipidated-analogs that exhibited slow homodimer dissociation rates resemble the original fragment hit with lipid-like moieties that vary in length (**Figure 2D**). We were interested in verifying that the kinetic behavior exhibited by the lipidated analogs in Figure 2C was due to binding in the homodimer central channel where we observe lipids binding.

> We attempted two strategies of crystallizing our lipidated-compounds with the apo-Orf9b homodimer to structurally resolve the complex. First, we attempted to soak our compounds into apo-Orf9b homodimer crystals but observed poor tolerance to DMSO which quickly disintegrated the crystals after minutes. Second, we attempted to cocrystallize the lipidated-compounds after refolding the Orf9b homodimer to generate apo-Orf9b homodimers but could not identify conditions that yielded crystals upon re-screening.

*As an alternative strategy,* we turned to a “co-folding” prediction model, Chai-1 (Discovery et al. 2024) to predict the structure of the Orf9b homodimer with the lipidated-analogs. All three analogs were predicted by Chai-1 to bind in the central channel as expected with reasonable ipTM scores for all three compounds (**Figure 2C**). We compared the Chai1 predicted structures to our original fragment bound structures which showed that the fragment portion of all three analogs were predicted to bind deeper inside the central channel than the original fragments. This suggested to us that while the lipidated analogs are predicted to bind in the central channel in support of our previous observations, their binding poses do not recapitulate the original lipid-fragment merged structures.

We next considered whether more than one lipidated analog was capable of binding to the central channel and therefore tested whether 2 analogs could bind to the channel at either end. For all three cases, Chai-1 predicted that two copies of the lipidated analogs could bind in the central channel (**Figure 2C**). In the case of compounds 1 and 3, placing two copies of the analog resulted in the fragment derived portion extending out of the central channel to resemble the original fragment-bound structure, albeit, with notable differences in the conformation of the fragment portion compared to the original structures. In the case of compound 2, the fragment portion of the analog is oriented in the opposite orientation of the original fragment whereas compound 1 is oriented in the same direction as the original fragment structure but shifted deeper into the channel by approximately 2Å. Compound 2, despite sharing the same parent fragment as compound 1 but with a lipid-moiety that is 2 carbons shorter, has its orientation flipped when two copies are used where the lipid tail extends out of the channel and the fragment portion is inserted deep into the channel (**Figure 2C**).

Incorporating restraints based on the experimental structure did not alter the predicted orientation of compound 2 when two copies are co-folded with Orf9b suggesting limitations in placing multiple ligand copies in the same binding site. None of these structures were publicly available on the PDB prior to these structural predictions, therefore, they were not available for Chai1 to train on. The placement of the lipidated-analogs in the central channel was in line with the anticipated binding mode, albeit, with discrepancies between the predicted and structurally observed portions of the analog compounds. Together, these results suggest that lipidated analogs can exert a stabilizing effect on the Orf9b homodimer and slow the rate of homodimer dissociation similar to the effect of co-purifying lipids, however, the exact structural basis for the observed kinetic changes remains to be more clearly defined.

### Developing a high-throughput fluorescence polarization screen to find Tom70-binding small molecules

As a complementary method to probing the mechanisms underlying Orf9b mediated suppression of IFN, we also sought to identify chemical probes that could bind to Tom70 and block Orf9b. We modified our fluorescence polarization peptide displacement assay for high-throughput screening of small molecule libraries to identify compounds that bound to Tom70 to block Orf9b binding (**Figure 3A**). In this assay, we generated peptides derived from the structurally resolved portion of Orf9b bound to Tom70 and appended a C-terminal fluorescein to create fluorescent peptide probes (**Figure 3–figure supplement 1**). When bound to Tom70, the complex emits a higher FP signal compared to the probe in solution on its own. We adapted a peptide fluorophore that we previously characterized by truncating several residues from the C-terminus to improve the signal to noise window which yielded a Orf9b peptide fluorophore with a Kd of 3.4uM and a ΔmP = 50mP which was a robust difference between bound and unbound fluorophore (Orf9bΔ10aa-FITC) (**Figure 3–figure supplement 1**). We next determined the sensitivity and reproducibility of our assay in a 384 well format which had a Z’ = 0.7 and a low variability in signal between wells which was suitable for a high-throughput screen.

**Figure 3:**
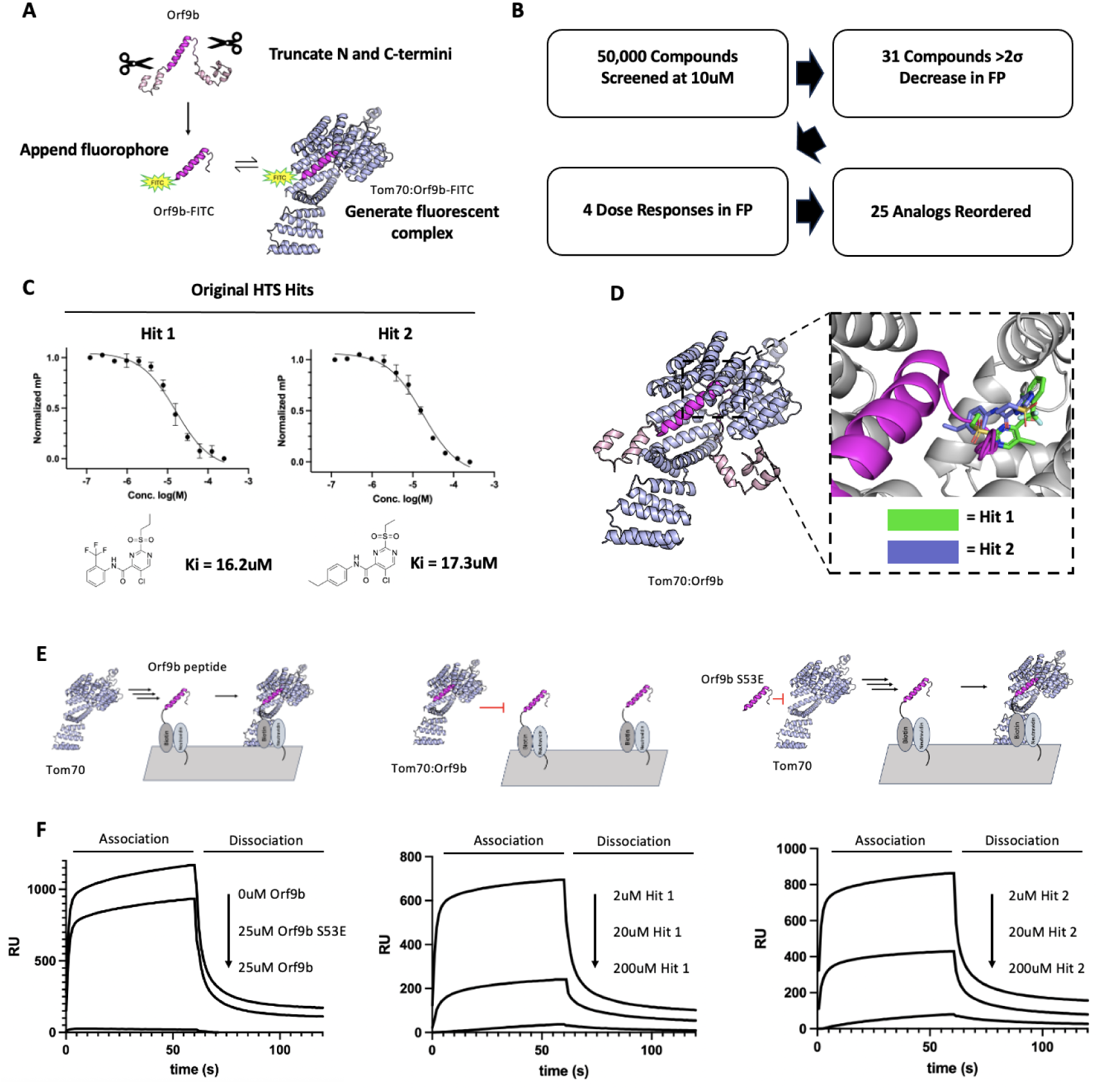
High-throughput screen identifies Tom70-binding small molecules that block Orf9b binding. (A) Diagram for generating fluorescent peptides derived from Orf9b for high throughput screening against Tom70. (B) Overview of high throughput screen and triage process. (C) Dose response curves for compounds identified by high throughput screen. (D) Chai-1 prediction of Tom70 co-folded with HTS hits (green and blue). Orf9b (magenta) is superimposed with compounds illustrating potential mechanisms for blocking Orf9b binding to Tom70. (E) Diagram of surface plasmon resonance experiments to measure binding of Tom70 to immobilized Orf9b peptide in the presence or absence of competitive binders derived from the Orf9b peptide. (F) SPR sensorgrams for Tom70 binding to Orf9b peptide immobilized surface in the presence or absence of competitive binders.

We screened 50,000 compounds from the Chembridge library at single concentrations of 10uM and selected 31 compounds that showed a 2 standard deviation decrease in FP signal for reordering after we filtered for fluorescent artifacts and individual plates that showed Z’ < 0.5. We reordered compounds and tested them at single concentrations of 200uM followed by dose responses of the top 3 compounds that showed the greatest decrease in FP signal (**Figure 3B**). 2 of those compounds showed Ki values in the double digit uM range with compounds we termed Hit 1 and Hit 2 both having the lowest Ki of 16.2uM and 17.3uM (**Figure 3B and C**). These compounds are highly similar to each other with the primary difference being the trifluoro group at the ortho position on hit 1 and the ethyl group at the para position on hit 2 (**Figure 3C**).

### Structure prediction tools suggest possible mechanisms of compound mediated inhibition of Orf9b binding

To investigate the structures of these compounds bound to Tom70, we used Chai-1 to co-fold Tom70 with the screening hits we identified. Both predictions had low ipTM scores of 0.5013 and 0.2639 and did not share the same orientation, however, both shared the same general binding location inside of the C-terminal domain of Tom70. When superimposing Orf9b, the predicted binding locations of both compounds could feasibly block Orf9b due to a steric clash with the N-terminal disordered region of Orf9b that extends back out of the C-terminal domain of Tom70 (**Figure 3D**). To further corroborate the Chai-1 predictions for the binding location of our screening hits to Tom70, we also used FTMap to explore prospective small-molecule binding sites on Tom70. FTMap analysis of Tom70 showed three main fragment clusters (**Figure 3–figure supplement 2**) within the C-terminal domain of Tom70 where Orf9b binds. Cluster 2 had the largest concentration of fragments bound which directly coincided with the predicted binding location of our screening hits to Tom70 lending further support to the predicted binding region on Tom70 (**Figure 3–figure supplement 2**). Therefore, even though the confidence scores are relatively low, the predicted complexes between our screening hits and Tom70, suggest that the mechanism that explains the inhibition of Orf9b binding is that our hits block part of the Tom70 C-terminal binding site.

### HTS hits block Tom70 from binding to Orf9b in an SPR format

To validate our screening hits, we next turned to surface plasmon resonance to determine if our hit compounds could inhibit Orf9b binding by competing for the Tom70 C-terminal binding site. We utilized a C-terminally biotin tagged Orf9b peptide based on the same amino acid sequence as the Orf9b-FITC fluorescent peptide which we immobilized to the surface of the chip. In this SPR format, when Tom70 is used as the analyte, we expect to see a robust increase in the sensorgram response as an indication of binding, however, if we pre-incubate Tom70 with an untagged Orf9b peptide, we should see little to no response due to saturation of the C-terminal binding site on Tom70 preventing Orf9b peptide on the chip surface from binding. Conversely, if we use a phosphomimetic Orf9b peptide, the phosphomimetic should be unable to bind to Tom70 leaving the C-terminal binding site free to bind to Orf9b immobilized on the surface (**Figure 3E**). We tested this SPR format and as expected observed a robust increase in the sensorgram response when Tom70 alone is used as the analyte. As a positive control, pre-incubating an excess of untagged Orf9b peptide with Tom70 resulted in very little response indicating that Tom70 was not binding to the surface due to obstruction of the C-terminal domain (**Figure 3F**). As a negative control, we pre-incubated an excess of the Orf9b S53E peptide with Tom70 which resulted in a restoration of the response indicating that the C-terminal domain on Tom70 was largely free (**Figure 3F**). We next tested our original HTS hits at three different concentrations to determine if they were bound to Tom70 by producing a decrease in the sensorgram response as compound concentration increased (**Figure 3F**). For both compounds, we observed a decrease in the measured response as the compound concentration increased indicating that these compounds were blocking Orf9b from binding to the C-terminal domain of Tom70 as we expected (**Figure 3F**).

### HTS analogs show improved binding affinity for Tom70

To improve upon our initial screening hits, we began exploring compound structure activity relationships (SAR) by testing how different changes to either the ortho, meta, or para positions affected compound affinity for Tom70. We ordered 25 analogs available through Enamine and determined the Ki for 22 compounds by 12-point dose response in the same FP assay format (**Figure 4–figure supplement 1**). We observed dynamic SAR across 22 analog compounds with 3 compounds showing no binding to Tom70 out to the highest concentration tested, 4 compounds showing weaker affinity than the original HTS hits, and 6 compounds showing no noticeable difference in affinity compared to the original HTS hits. We identified several compounds with improved single digit micromolar affinity. Compounds 21 and 22 showed approximately 3-fold improvement in affinity compared to the original HTS hits. Given the similarity between our analog compounds and the original screening hits, we wanted to determine if the predicted binding pose and location to Tom70 was consistent with the predicted binding of the original screening hits.

**Figure 4:**
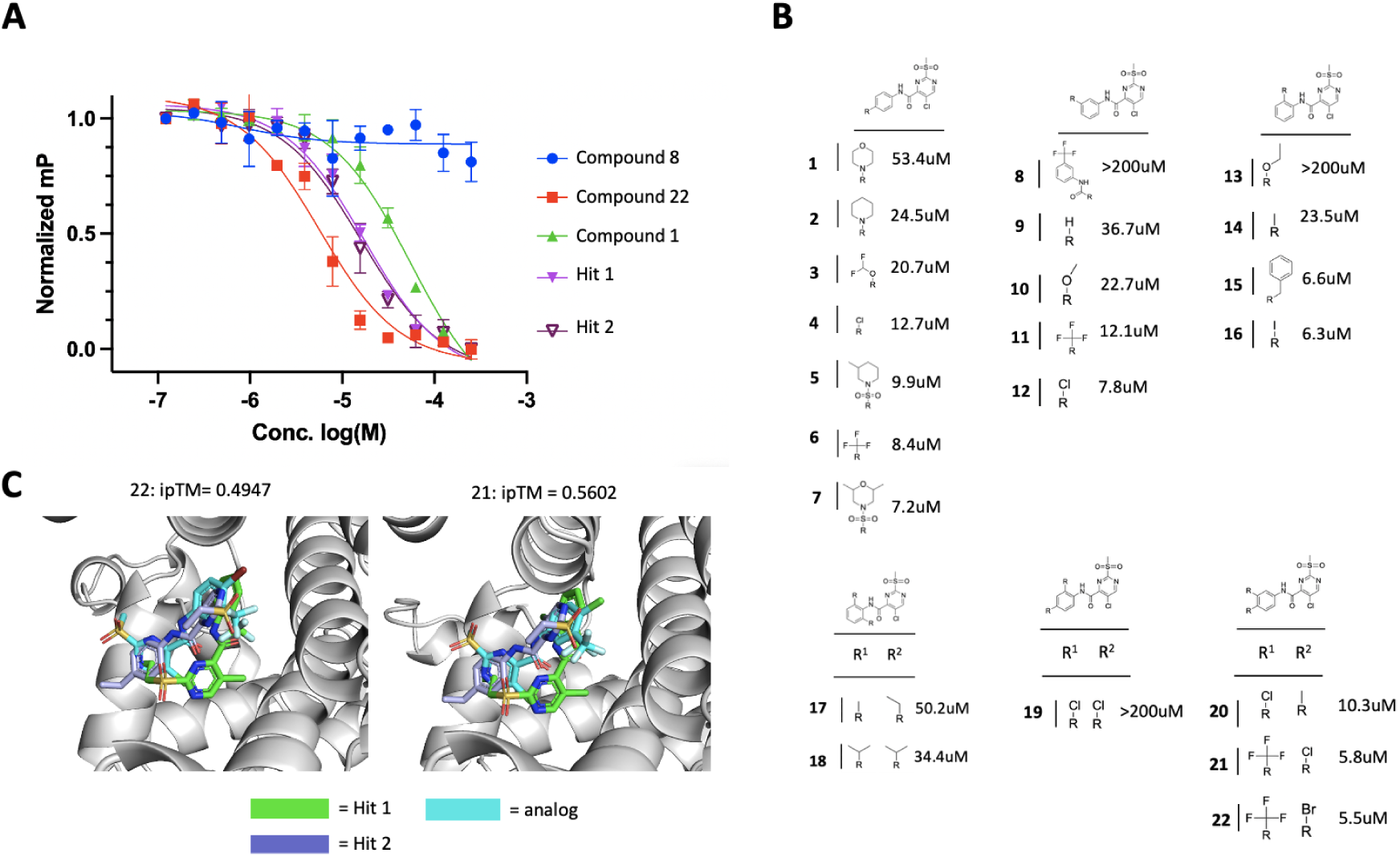
Summary of Tom70-binding analog compounds. (A) Dose response of select HTS analog compounds in FP format. (B) Table of purchased analogs showing the relationship between changes to the benzene group and affinity for Tom70. Compounds that had no measurable effect on FP were reported to have Ki >200uM. (C) Chai-1 predictions of Tom70 co-folded with top binding analogs with ipTM scores. Original HTS hits co-folded with Tom70 are shown in blue and green and analog compounds are shown in cyan.

We again used Chai-1 to co-fold Tom70 with all 23 analogs and observed that all compounds bound within roughly the same location of Tom70 but had low predicted ipTM scores no better than 0.5602 with most compounds scoring between 0.2 and 0.3, similar to our original screening hit predictions indicating low confidence in placement. In the cases of compounds 21 and 22, both compounds showed consistency in the predicted binding location compared to the original HTS hits and produced the highest ipTM scores relative to all other compounds predicted. Compared to the predicted binding poses and locations of the original screening hits, all 4 compounds showed a similar binding pose, however, these predictions can only give us an idea about where they may bind and not the specific interactions they make with Tom70 for aiding SAR.

## Discussion

Orf9b is a fold-switching antagonist to innate immunity when bound to the host mitochondrial receptor Tom70 (Meier et al. 2006; San Felipe et al. 2025; Jiang et al. 2020; Gordon et al. 2020). Inhibition of Orf9b binding to Tom70 has been shown to restore interferon signaling by mutagenesis that mimics a phosphorylation blocking the direct interaction (Thorne et al. 2022), however, there are currently no known small molecules that inhibit Orf9b’s function. We identified two potential strategies for targeting Orf9b and Tom70 with small molecules to inhibit this interaction using both high throughput and crystallographic fragment screens. We identified two biologically distinct sites for fragment binding on the Orf9b homodimer using X-ray crystallography: an external homodimer interface composed of the two chains of Orf9b and the entrance to the lipid-binding central channel. We explored both sites starting from our initial fragment hits to identify analog compounds that could stabilize Orf9b in the homodimer conformation. Despite most of our observed fragments binding at an external interface composed of the two chains of Orf9b, analog compounds bound weakly and showed no noticeable stabilization of the homodimer. However, the addition of lipid-like moieties to fragments that bound near the central channel entrance showed a noticeable increase in homodimer stability compared to the apo-homodimer by slowing the dissociation of the homodimer into monomers, similar to the effect exerted by lipid-binding. While we do not have a specific PEG/lipid control that recapitulates the behavior of lipid-bound Orf9b to directly compare with these analogs, the fact that only some analogs slowed the dissociation indicates that this is a specific effect of the small molecule head group attached to the lipid tail. Therefore, lipid-binding central channel may be a more suitable target to exploit with a small molecule to inhibit Tom70 binding through Orf9b homodimer stabilization.

Complementary to our crystallographic approach focused on the Orf9b homodimer, we also conducted a traditional high-throughput screen (HTS) to identify compounds bound to Tom70 to inhibit Orf9b monomer binding. Our HTS identified several hits with two closely related compounds showing low double digit micromolar affinity for Tom70. While we were unable to obtain experimental structures of these compounds bound to Tom70 to pinpoint the exact binding location, co-folding predictions suggested that these compounds could be binding deep within Tom70’s C-terminal domain. When superimposed with the experimentally resolved structure of Orf9b bound to Tom70, the compound competes for the same surface and would have steric clashes that block Orf9b binding. Limited analoging of these HTS hits yielded compounds with a 3-fold improvement in affinity for Tom70 in the single digit micromolar range. Although these compounds are relatively weak, they serve as tools for further probing the Orf9b-Tom70 interaction. *Although we have not assessed the permeability of these compounds and their ability to restore IFN in the context of cell based assays, one way this could be accomplished is by adapting a previously used reporter cell line (San Felipe et al 2025). Specifically, this assay uses a reporter cell line that has a Lucia reporter under the control of the IFN-β/ISG56 promoter that is expressed upon mock infection with synthetic dsRNA. When WT Orf9b is co-transfected with the reporter, IFN is suppressed, and, we have shown that phosphomimetic mutants of Orf9b that block its interaction with Tom70 restores IFN signaling. If these compounds are sufficiently permeable, then, for the lipidated-compounds targeting the Orf9b homodimer, we would expect that if the Orf9b monomer:homodimer equilibrium would shift towards the homodimer due to the stabilizing effect these lipidated-compounds have on the homodimer conformation leading to an increase in IFN signaling. Similarly, for compounds targeting Tom70, we would expect that these compounds block the Orf9b monomer from binding to Tom70 to produce a similar effect.* In the future, we hope these tools can be used to probe the dual role of Tom70 in facilitating mitochondrial import and innate immune activation.

Collectively, the results from our two screens demonstrate that it is possible to develop chemical matter against both Orf9b and Tom70 which can be used to inform on the mechanisms underlying Tom70-mediated innate immune activation. Our results expand on the targetable space for both Orf9b and Tom70. Our work will serve as chemical starting points to explore both the Tom70-Orf9b equilibrium during coronavirus infection as well as the mechanisms underlying innate immune activation.

## Methods and Materials

### Orf9b purification

The WT Orf9b gene was cloned into a pET-29a backbone with an N-terminal 6xHis tag and TEV protease cleavage site. BL21 E. coli cells from NEB were transformed and grown in a starter culture of Luria Broth at 37C overnight. 1L cultures were inoculated with 10mL of the starter culture and grown in Terrific Broth media supplemented with Kanamycin (100ug/mL) at 37C until an optical density of 0.6-0.8 was reached. Cultures were chilled at 4C for 15 minutes and then induced with 1mM IPTG and grown at 16C overnight. Frozen cell pellets were resuspended in a lysis buffer of 300mM NaCl, 50mM Tris-HCl pH 7.5, and 0.5mM TCEP and a cOMPLETE EDTA free protease inhibitor tablet added per liter followed by sonication. Lysate was clarified by centrifugation and loaded onto a 5mL HisTrap HP Ni-NTA (Cytiva: 17524802) column pre-equilibrated with the lysis buffer supplemented with 10mM Imidazole. The tagged protein was eluted off the column with a buffer of 300mM NaCl, 50mM Tris-HCl pH 7.5, 300mM imidazole, and 0.5mM TCEP. The 6xHis tag was cleaved by TEV protease protease added in a ratio of 1:20 by mass and dialyzed into a buffer of 150mM NaCl, 50mM Tris-HCL pH 7.5, 5% glycerol, and 0.5mM TCEP overnight. The cleaved protein was separated by rerunning the cleaving reaction over a Ni-NTA column and collecting the flow through. The cleaved protein was then concentrated with a 3kDa Amicon centrifugal filter and purified by size exclusion chromatography using an Superdex 75 16/600 column equilibrated with 100mM NaCl, 20mM HEPES pH 7.5, and 5% glycerol. Eluted fractions were pooled, filtered and flash frozen in liquid nitrogen, and stored at −80C.

### Refolded Orf9b purification

WT Orf9b was cloned into a pET-29a backbone with an N-terminal 6xHis tag and TEV protease cleavage site. BL21 E. coli cells from NEB were transformed and grown in a starter culture of Luria Broth at 37C overnight. 1L cultures were inoculated with 10mL of the starter culture and grown in Terrific Broth media supplemented with Kanamycin (100ug/mL) at 37C until an optical density of 0.6-0.8 was reached. Cultures were then induced with 1mM IPTG and grown at 16C overnight. Frozen cell pellets were resuspended in a lysis buffer of 300mM NaCl, 50mM Tris-HCl pH 7.5, and a cOMPLETE EDTA free protease inhibitor tablet added per liter followed by sonication. Lysate was clarified by centrifugation and loaded onto a 5mL HisTrap HP Ni-NTA column pre-equilibrated with the lysis buffer supplemented with 10mM Imidazole. 10 column volumes of buffer (6M Guanidine HCl, 300mM NaCl, 50mM Tris-HCl pH 7.5, and 0.1%Triton X-1000, 0.5mm TCEP) was flowed and the tagged protein was eluted with 6M Guanidine HCl, 300mM NaCl, 50mM Tris-HCl pH 7.5, 300mM imidazole, and 0.5mM TCEP. The denatured tagged Orf9b was diluted to less than 65ug/mL in a refolding buffer composed of 550mM Guanidine HCl, 55mM Tris, 21mM NaCl, 0.88mM KCl at pH 8.2 overnight at 4C and allowed to refold. The refolded 6xHis tag was cleaved by TEV protease, added in a ratio of 1:20 by mass and dialyzed into a buffer of 150mM NaCl, 50mM Tris-HCL pH 7.5, and 0.5mM TCEP overnight. The cleaved protein was separated by rerunning the cleaving reaction over a Ni-NTA column and collecting the flow through. The cleaved protein was then concentrated with a 3kDa Amicon centrifugal filter and purified by size exclusion chromatography using an Superdex 75 16/600 column equilibrated with 100mM NaCl and 20mM HEPES pH 7.5 and 5% glycerol.

### Biotinylated Orf9b purification

The WT Orf9b plasmid was used for generating biotinylated Orf9b for SPR experiments. An AVI-tag was introduced to the N-terminus using a Q5 Site-Directed Mutagenesis kit (NEB: E0544S) and confirmed by sequencing. BL21 cells were cotransfected with the AVI-tagged plasmid and BirA biotin ligase plasmid in equal quantities by mass. WT Orf9b AVI-tagged plasmid was kanamycin resistant and BirA was chloramphenicol resistant for double selection. Starter cultures were grown ON at 37C in LB media. 1L cultures were inoculated with 10mL of the starter culture and grown in Terrific Broth media supplemented with both chloramphenicol and kanamycin (100ug/mL) and grown at 37C until an optical density of 0.6-0.8 was reached. 1L cultures were then induced with 1mM IPTG and 100uM biotin and grown overnight at 16C. Purification proceeded as previously described with the modification of not including a TEV cleavage reaction.

### Tom70 purification

hTom70 (109-600) lacking the transmembrane domain was cloned into a pET-29b(+) backbone with an N-terminal 6xHis tag and SUMO tag. Transformation and growing conditions were the same as WT Orf9b other than the media being supplemented with Carbanicillin (100ug/mL). HisTrap HP Ni-NTA column purification conditions and buffers were also the same as Orf9b except for the cleavage of the 6x-His and SUMO tag which was performed with Ulp1 protease at a ratio of 1:20 by mass. All other conditions were the same except for size exclusion chromatography which was performed with a Superdex 200 16/600 column.

### Orf9b crystallization, Fragment soaking, and Data Collection

WT Orf9b was concentrated to 10mg/ml and crystallized by sitting drop diffusion with a reservoir buffer of 10% PEG3350, 10% PEG1000, 10% MPD, 0.15M Ethylene Glycol, 0.1M MES pH 6.5, 0.1M Imidazole pH 6.5 at 20C. Crystallization drops were set up with 200nL of Orf9b mixed with 100nL of the mother liquor and 100nL of 1:10,000 Orf9b microcrystals in SWISSCI 3-well plates.

Crystals appeared 1 day after plating and grew to a maximum size in 7 days. All crystals appeared 1 day after plating and grew to a maximum size in 7 days. Fragment libraries were dispensed into crystal drops using an Echo650 acoustic liquid handler. 320 fragments from the Enamine essential and 91 fragments from the UCSF_91 libraries were soaked into Orf9b crystals. 100nL of fragment solutions were soaked into 400nL crystal drops. Plates were resealed and allowed to sit at 20C for 24 hours. To generate background datasets for PanDDA analysis, 19 crystals were soaked with 100nL of DMSO for 24 hours at 20C. Crystals were looped and flash frozen without added cryo-protectant. X-ray diffraction data was collected at the Advanced Light Source beamline 8.3.1.

### Data Refinement and Modeling

Diffraction data was indexed in XDS (Kabsch 2010) and merged and scaled using Aimless (Evans and Murshudov 2013). A single high-resolution DMSO soaked dataset was used for generating R-free flags which were copied over into each DMSO and fragment soaked dataset. Molecular replacement was performed with Phaser (Winn et al. 2011) and initial refinement was performed using Refmac (Vagin et al. 2004) using the DIMPLE pipeline using a high-quality DMSO soaked structure of the Orf9b homodimer as reference. Fragment analysis was performed using PanDDA (Pearce et al. 2017b) using 19 DMSO soaked datasets for generating background electron density maps with the highest resolution and lowest R_free_ values. PanDDA was run with default settings and fragments were modeled into PanDDA event maps using Coot (Emsley et al. 2010). Fragment restraints were generated using phenix.elbow (Moriarty et al. 2009) using SMILES strings. Group deposition PDB codes were assigned as follows:

13TC (x016), 13TD (x026), 13TE (x027), 13TF (x030), 13TG (x035), 13TH (x036), 13TI (x040), 13TJ (x043), 13TK (x044), 13TL (x057), 13TM (x063), 13TN (x116), 13TO (x120), 13TP (x159), 13TQ (x166), 13TR (x169), 13TS (x176), 13TT (x184), 13TU (x228), 13TV (x234), 13TW (x247), 13TX (x259), 13TY (x263), 13TZ (x267), 13UA (x270), 13UB (x271), 13UC (x279), 13UD (x281), 13UE (x287), 13UF (x291), 13UG (x298), 13UH (x302), 13UI (x304), 13UJ (x312), 13UK (x314), 13UL (x317), 13UM (x321), 13UN (x322), 13UO (x336), 13UP (x342), 13UQ (x347), 13UR (x355), 13US (x399), 13UT (x407), 13UU (x409), 13UV (x412), 13UW (x417), 13UX (x420), 13UY (x432), 13UZ (x434), 13VA (x436)

Statistics for all deposited structures listed can be found in Supplementary File 1. PanDDA event and Z-maps used for modeling fragments are available at: https://doi.org/10.5281/zenodo.19750677

### Fluorescence Polarization High-Throughput Screen

To generate the fluorescent complex, a master mix of 2.5uM Tom70 and 50nM Orf9bΔ10aa-FITC were incubated together in a buffer of 100mM NaCl, 20mM HEPES pH 7.5, 0.05% Tween-20, and 0.1mM TCEP for 1 hour. The master mix was dispensed into 384 well plates (Corning: 4514) using a Multidrop Combi Reagent Dispenser at 20uL per well. Control wells used for calculating Z’-value were either columns 1-2 with 10nL of DMSO as a negative control or columns 23-24 with 10uM of Orf9b peptide as a positive control. Control wells were aliquoted using an Echo650 acoustic liquid dispenser. Columns 3-22 were aliquoted with 10nL of compound from the Chembridge Diversity Library (50,000 compounds) at a final compound concentration of 10uM using an Echo650 acoustic liquid dispenser and allowed to equilibrate with the fluorescent complex for 4 hours at room temperature. Plates were read using an Envision HTS Dual mode plate reader measuring fluorescence polarization (excitation: 485nm, emission: 535nm). Hits were selected that showed greater than 2 standard deviation decreases in FP for reordering. Competitor binding curves were generated in 12-point duplicates using serial 2-fold dilutions. Ki values were calculated in GraphPad PRISM using a one-site binding model and the Kd of the Tom70:Orf9bΔ10aa-FITC complex.

### Surface Plasmon Resonance

For measuring the interaction between Orf9b and Tom70 in the presence of HTS compounds, biotinylated Orf9b core peptide (44-70) was synthesized by Biomatik biotinylated at the C-terminus of the peptide. The Orf9b peptide was immobilized to a CMD500M chip (Xan-tec: SPSMCMD500M). 250nM of biotinylated Orf9b core peptide was immobilized for 60s at 25uL/s to either spots B or C on a neutravidin coated surface followed by 120s of dissociation. Spot A on the chip was used as a reference surface and had no Orf9b peptide immobilized to it. 2.5uM Tom70 was incubated with either Orf9b core peptide or Orf9b core peptide (S53E) at 25uM each, or HTS compounds in concentrations ranging from 2-200uM. All SPR experiments were performed using a running buffer of 100mM NaCl, 20mM HEPES pH 7.5, 0.5mM TCEP, 5% DMSO, and 0.05% Tween20. For measuring fragment analogs binding to Orf9b homodimer, biotinylated Orf9b homodimer was immobilized to a CMD500M chip for 120s at 25uL/s to either spots B or C on a neutravidin coated surface followed by 120s of dissociation. Analog compounds were resuspended in 100% DMSO then serially diluted into running buffer to achieve a final buffer composition of 100mM NaCl, 20mM HEPES pH 7.5, 0.5mM TCEP, 5% DMSO, and 0.05% Tween20. To account for the high refractive index of DMSO, a calibration curve was generated using DMSO ranging from 1-5%. Blank injections of the running buffer were regularly interspersed between analyte injections. Binding measurements were performed using a Bruker SPR-24 Pro. All SPR experiments were performed using a running buffer of 100mM NaCl, 20mM HEPES pH 7.5, 0.5mM TCEP, 5% DMSO, and 0.05% Tween20. All binding sensorgrams were double reference subtracted from the reference surface and the running buffer blank injections.

### Fluorescence Polarization Kinetic Assay

Orf9b core peptide was synthesized by Biomatik comprising residues 44-70 of Orf9b (core peptide.) Orf9bΔ10aa-FITC was synthesized comprising residues 44-60 with a C-terminal fluorescein fluorophore. All peptides were resuspended from lyophilized powder in a 3:1 ratio of 100mM NaCl, 20mM HEPES pH 7.5 and DMSO and filtered. The Tom70:Orf9bΔ10aa-FITC fluorescent complex was generated by mixing with 2.5uM Tom70 in a buffer of 100mM NaCl, 20mM HEPES pH 7.5, 0.5mM TCEP with 200nM of Orf9bΔ10aa-FITC in black eppendorf tubes and equilibrated for 1hr. The fluorescent complex was added to black 96 well plates with a non binding surface (Greiner: 655900). Orf9b analog compounds were resuspended in DMSO and mixed with refolded WT Orf9b homodimer to a 10x working stock, the working stock was then chased into the preequilibrated wells containing the fluorescent complex to reach a final 1x concentration to initiate kinetic experiments. A solution of 20nM Orf9bΔ10aa-FITC in assay buffer was used to determine the background FP signal. Control experiments to assess if analog compounds acted on the Orf9b monomer were performed using the Orf9b core peptide as a proxy. Measurements were performed on a Tecan Spark running Magellan with a monochromator (excitation: 485nm, emission: 535nm). Measurements were taken in 15 second intervals between measurements with 7 seconds of shaking between intervals. Kinetic data was modeled using Berkeley Madonna v10.6.1 (Marcoline et al. 2022) as previously described (San Felipe et al 2025) to estimate Orf9b homodimer dissociation rates in the presence of analog compounds.

### Chai1 Co-Folding Prediction

Chai1’s web server was used for co-folding either Orf9b or Tom70 with their respective hits. For the Orf9b homodimer, the sequence for a single chain was supplied to represent chains A and B in the homodimer and the ligands were supplied as SMILEs strings. In the case where two copies of the ligand were investigated, the number of copies of the ligand was increased to two. Co-folding was performed using MMSeqs2 for the MSA and template searching to ensure the correct folding of the homodimer conformation. Tom70 co-folding was performed by supplying the sequence for hTom70 as a single chain and the compounds as SMILEs strings. Co-folding was performed using MMSeqs2 for the MSA with template searching.

## Acknowledgements

We thank Mark Herzik, Qiaozhen Ye, Kevin Corbett, Maxwell Bachochin for helpful discussions on potential Tom70 ligand binding modes. We acknowledge Jason Gestwicki and John Gross for helpful feedback, Amanda Paulson for data management and Jezrael Lafuente Revalde for equipment and dispensing support within the UCSF Small Molecule Discovery Center. This work was funded by NIH U19AI171110 (to NJK) and GM145238 (to JSF). The diffraction data of structures reported in this work was collected at beamline 8.3.1. of the Advanced Light Source (ALS). The ALS, a U.S. DOE Office of Science User Facility under contract no. DE-AC02-05CH11231, is supported in part by the ALS-ENABLE program funded by the NIH, National Institute of General Medical Sciences, grant P30GM124169.

## Competing Interests

N.J.K. has monetary and/or stock compensation with the following companies: GEn1E Life-sciences, Maze Therapeutics, Mreza Therapeutics, Rezo Therapeutics, and Tenaya Therapeutics. J.S.F.: J.S.F. holds equity in Relay Therapeutics, Impossible Foods, Arda Therapeutics, Profluent Bio, Interdict Bio (co-founder), Vilya Therapeutics, and Edison Scientific Inc and is a paid consultant for Relay Therapeutics, Profluent Bio, Vilya Therapeutics, and Monimoi Therapeutics. All other authors declare no competing interests.

---

**Figure 1–figure supplement 1:**
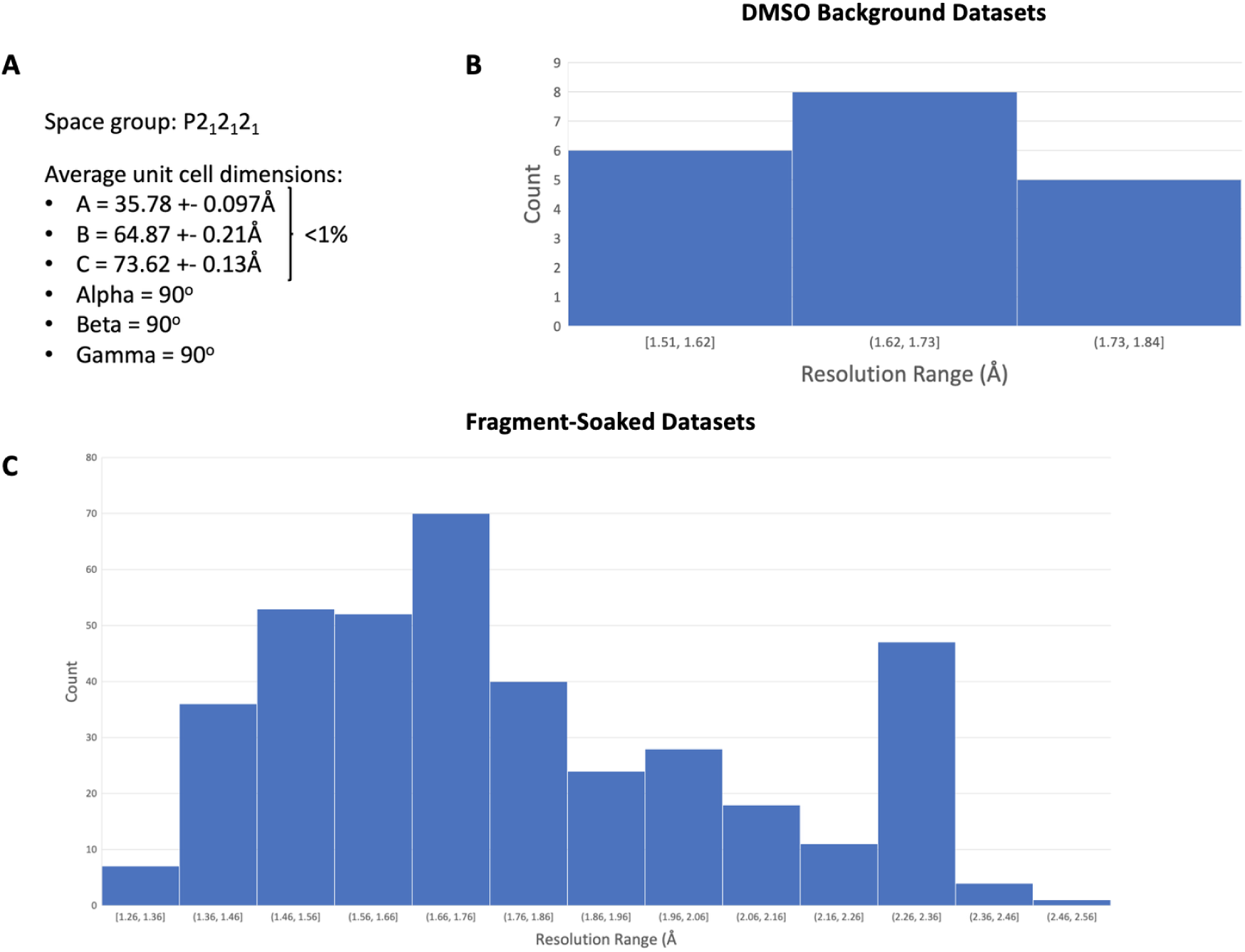
Summary of DMSO and fragment soaked data sets. (A) Orf9b homodimer crystal system unit cell parameters. All datasets collected showed less than 1% variation in unit cell dimensions. (B) Distribution of resolution ranges collected for DMSO soaked crystals used for PanDDA background maps. (C) Distribution of resolution ranges for fragment soaked datasets collected for conducting PanDDA analysis.

**Figure 1–figure supplement 2:**
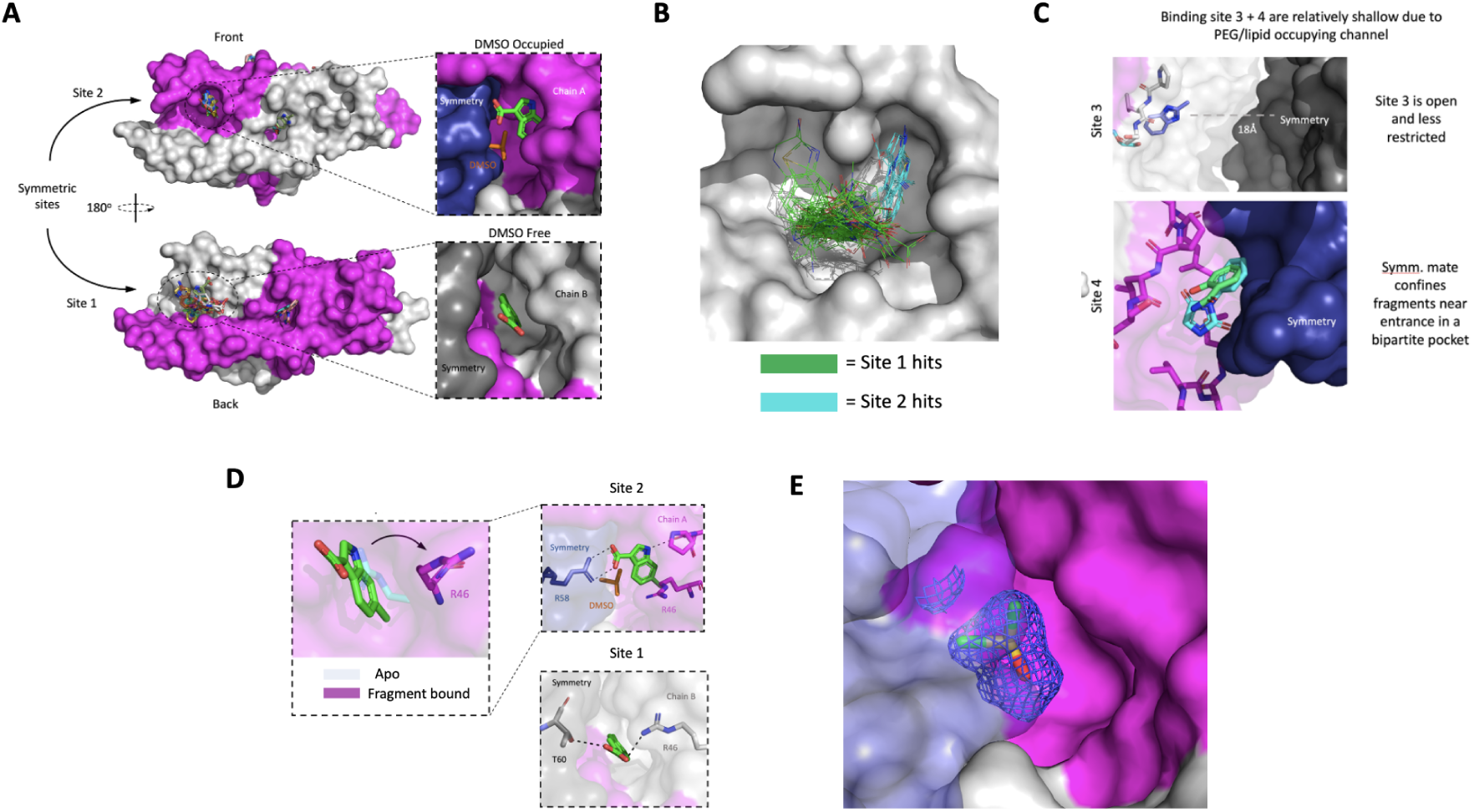
Comparison of fragment binding poses in sites 1 and 2. (A) Comparison of fragment binding sites 1 and 2. Binding site 2 is partially occupied by a DMSO molecule and has fragments that are capable of forming favorable hydrogen bonds with the neighboring symmetry mate. (B) Superimposition of all modeled fragments from sites 1 onto site 2. Fragments at site 2 (cyan) bind in a region and orientation in the binding site that is distinct to site 1 fragments (green) (C) Comparison of neighboring symmetry mates at fragment binding sites 3 and 4. Binding site 3 fragments are approximately 22A away from the nearest symmetry mate whereas site 4 fragments are in close proximity to a symmetry mate near the central channel entrance. (D) Observed conformational change of R47 on chain B upon fragment binding. Differences in fragment binding poses observed in sites 1 and 2 with neighboring symmetry mate. (E) Electron density modeled as DMSO occupies part of binding site 2 reducing pocket size for fragment binding.

**Figure 2–figure supplement 1:**
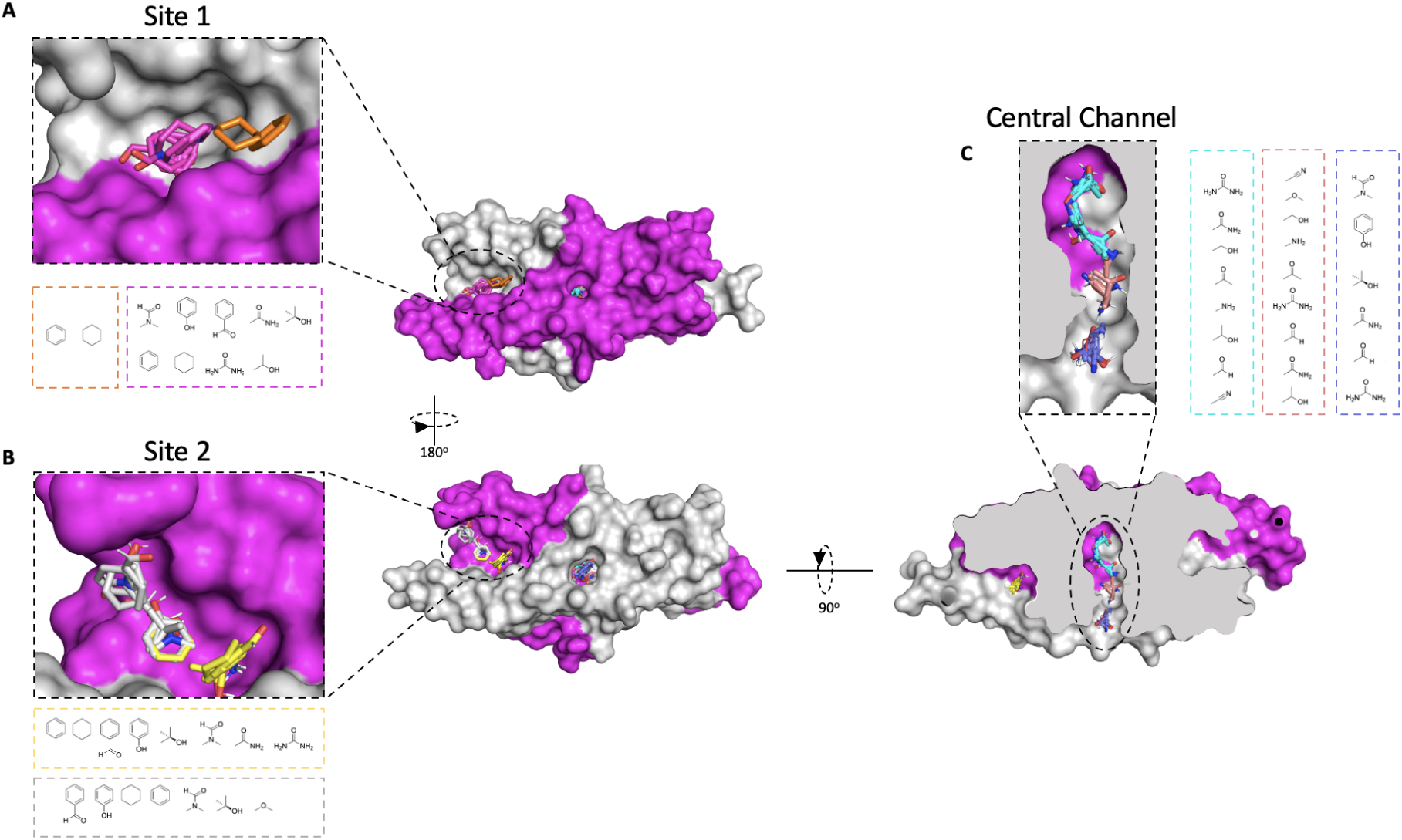
Overview of FTMap predicted fragment hot-spots. (A) Distribution of FTMap probes bound at site 1 on the Orf9b homodimer. (B) Distribution of FTMap probes bound at site 2 on the Orf9b homodimer. (C) Distribution of FTMap probes bound to the central channel of the Orf9b homodimer.

**Figure 2–figure supplement 2:**
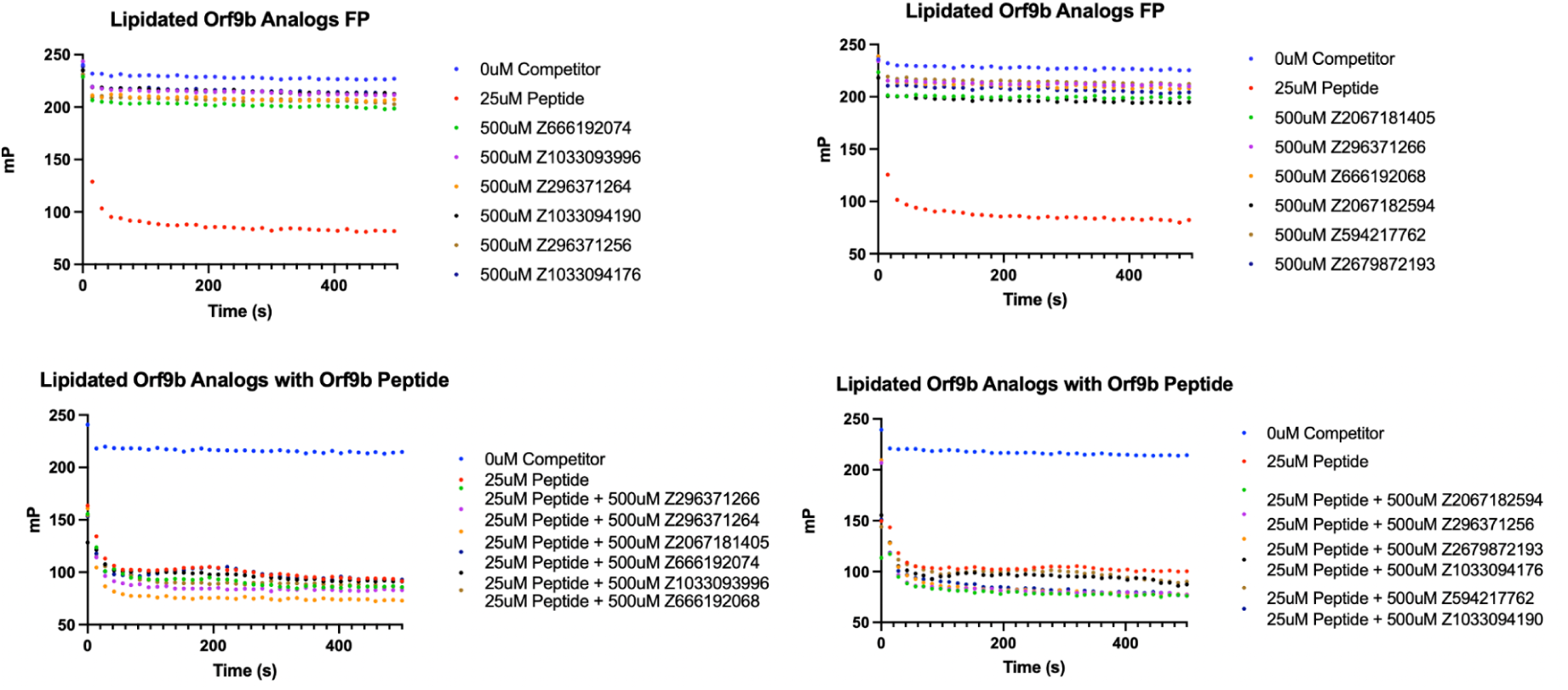
Lipidated analogs do not act on monomeric Orf9b or Tom70 directly. Lipidated Orf9b analogs do not compete with the fluorescent probe for binding to Tom70 in kinetic assay format. Control peptide exhibits a sharp decrease in FP signal over time as a positive control.

**Figure 3–figure supplement 1:**
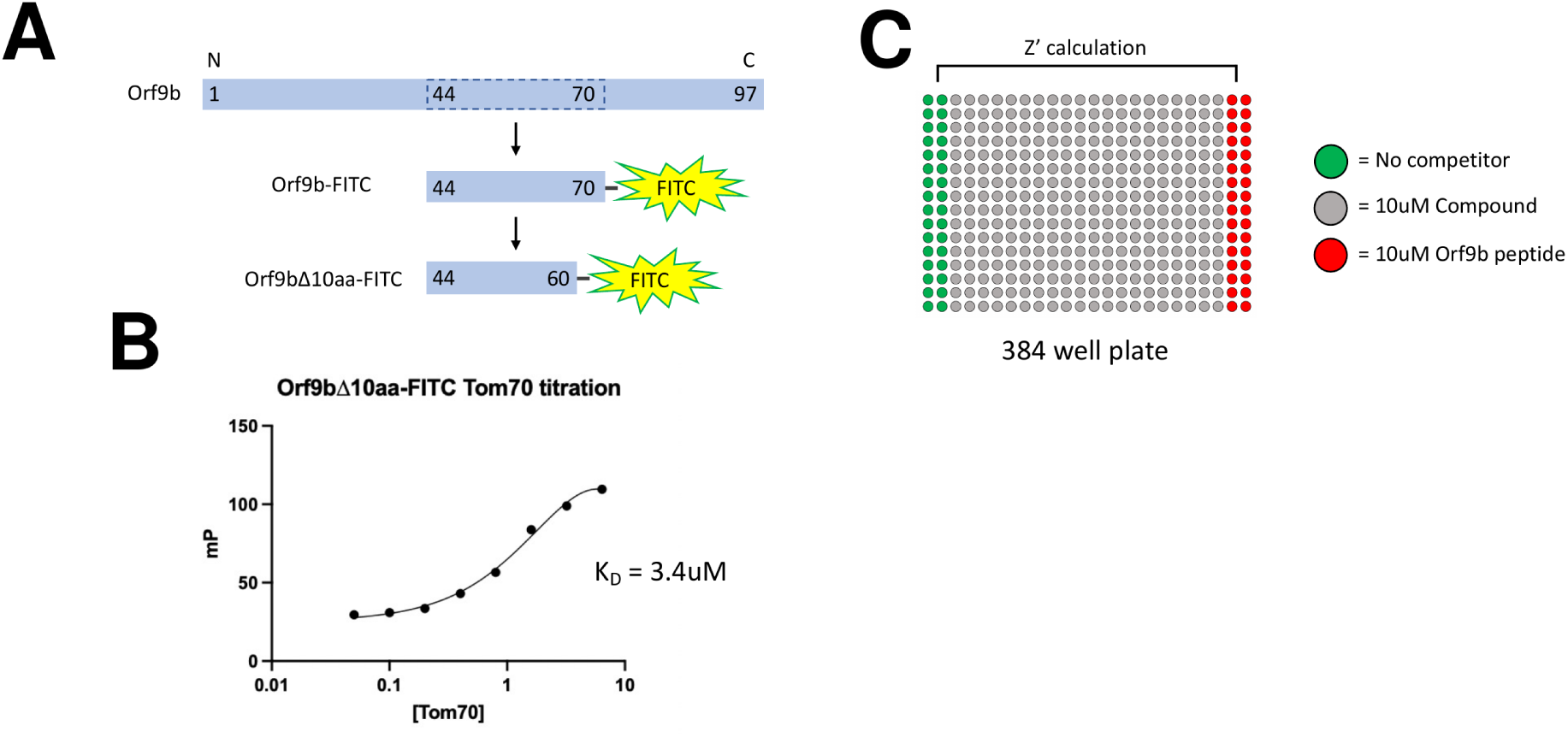
Design of fluorescent Orf9b peptides and HTS screen. (A) Design of fluorescent peptides derived from WT Orf9b. The structurally resolved residues of Orf9b bound to Tom70 (44-70) were used for generating C-terminally appended fluorescein peptides. A further 10 amino acids are truncated from the C-terminus of the Orf9b-FITC construct to generate the Orf9bΔ10aa-FITC fluorescent peptide used in the high throughput screen. (B) Titration of Tom70 against a fixed concentration of Orf9bΔ10aa-FITC was performed to identify the Kd. A non-linear regression single binding site model was used to determine the Kd of 3.4uM (C) Overview of a high-throughput screen set up. HTS was performed in 384 well plates with all wells containing the Tom70:Orf9bΔ10aa-FITC complex. Columns 1-2 contain only DMSO added and columns 23-24 contain 10uM of the Orf9b peptide as a positive control. Z’ values were calculated from columns 1-2 and 23-24.

**Figure 3–figure supplement 2:**
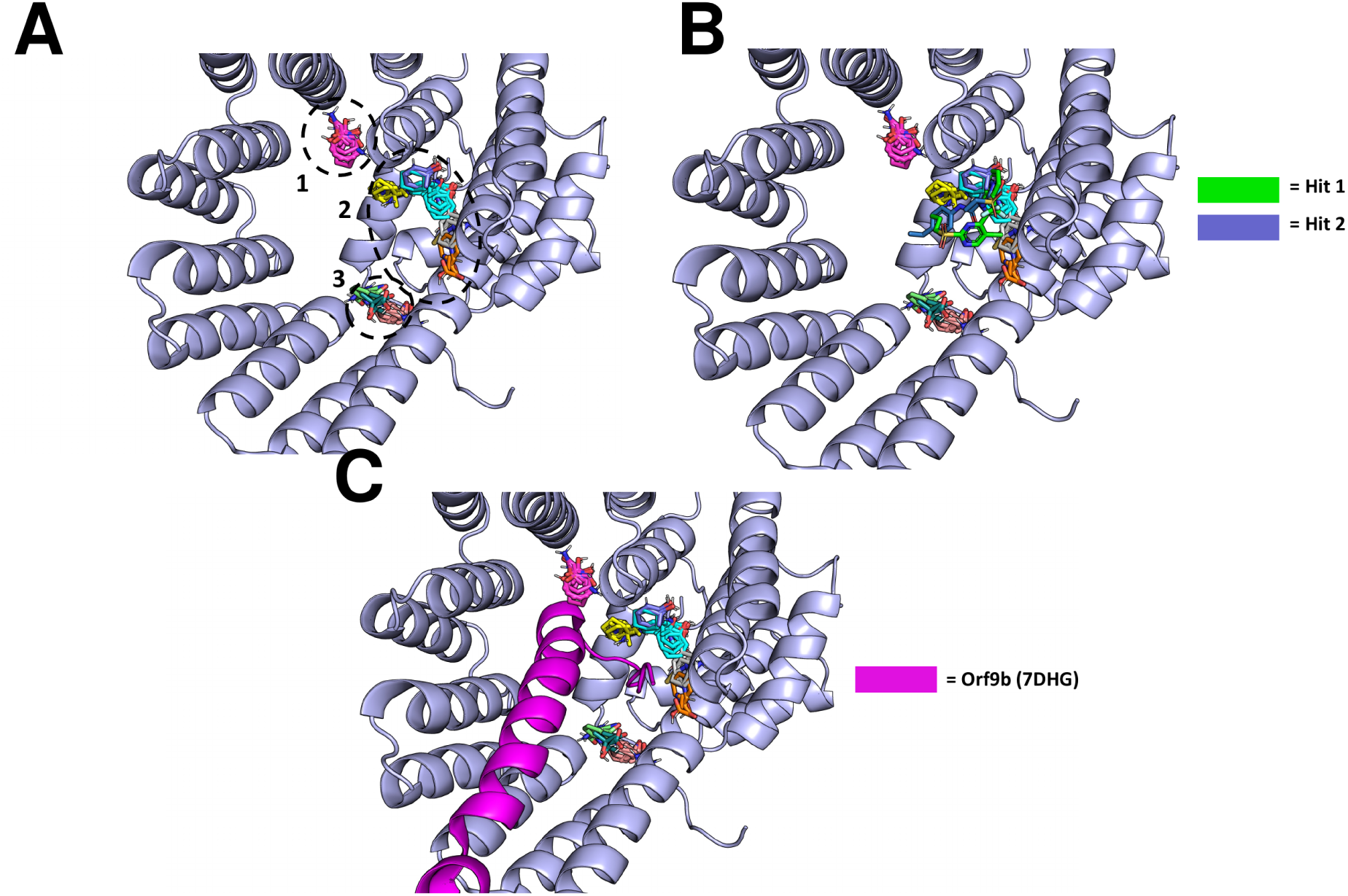
Structure predictions for Tom70-small molecule binding sites. (A) FTMap probe clusters within the Tom70 C-terminal binding domain. Clusters are grouped and number 1-3. (B) Superimposition of Chai-1 predicted binding locations for HTS Hits 1 and 2 with FTMap probe clusters. (C) Superimposition of Chai-1 predicted binding locations for HTS Hits 1 and 2 with FTMap probe clusters and Orf9b from the Cyro-EM structure of the Orf9b:Tom70 complex (PDB 7DHG).

**Figure 4–figure supplement 1:**
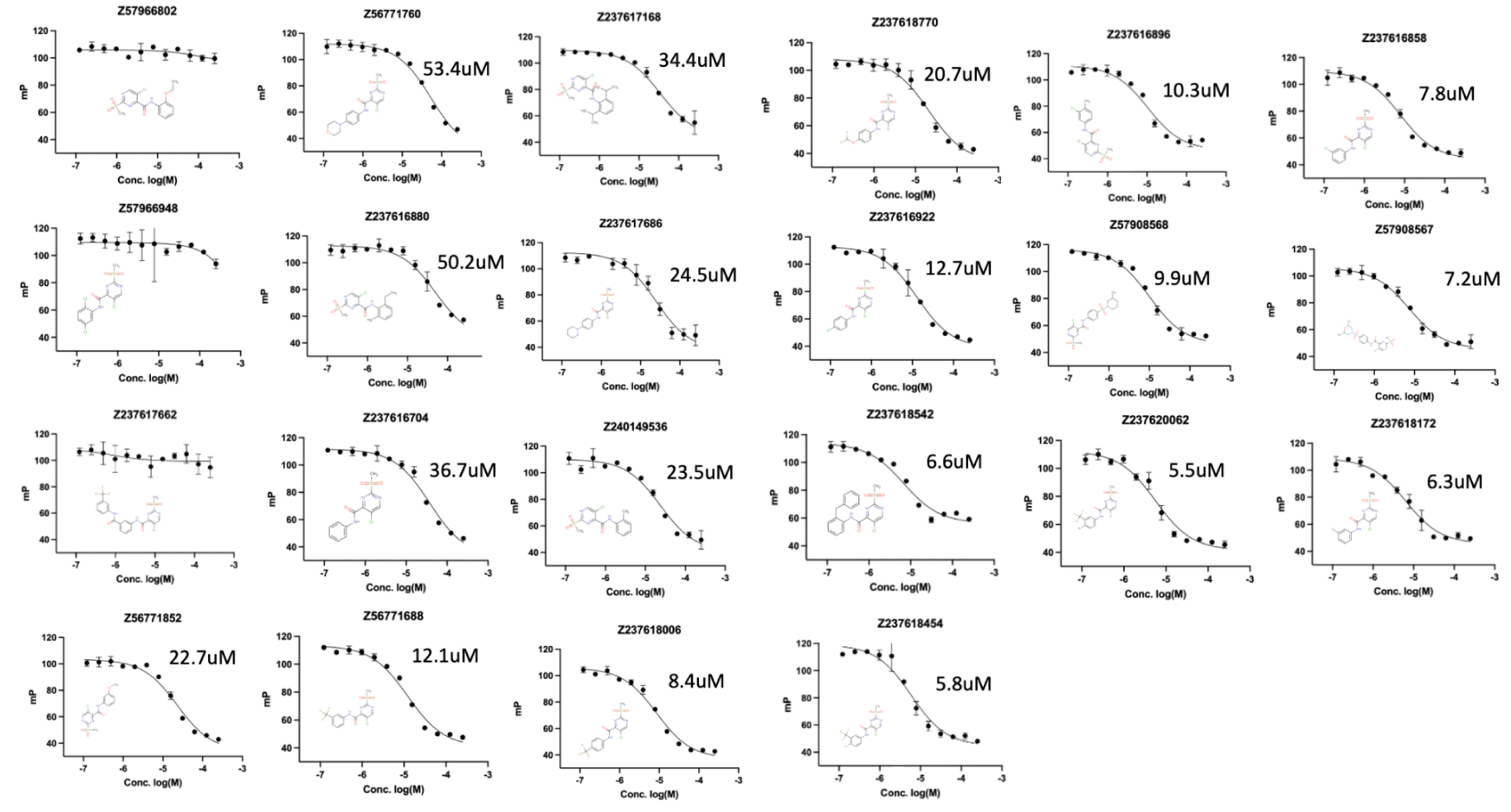
Dissociation Constants and Competition Binding Curves of Tom70 Analog Compounds. Competition binding curves of all tested Tom70 analog compounds in FP format. Error bars are for duplicate measurements with fitted curves shown using a one site fit for calculating Ki values. Structures of the compounds tested are shown in the bottom left.

## Supplementary Information

**Supplementary File 1.** Crystallographic data collection and refinement statistics for all deposited Orf9b structures (PDB 13TC-13VA).

## Notes

https://zenodo.org/records/19750678

### Competing Interest Statement

N.J.K. has monetary and/or stock compensation with the following companies: GEn1E Lifesciences, Maze Therapeutics, Mreza Therapeutics, Rezo Therapeutics, and Tenaya Therapeutics. J.S.F. holds equity in Relay Therapeutics, Impossible Foods, Arda Therapeutics, Profluent Bio, Interdict Bio (co-founder), Vilya Therapeutics, and Edison Scientific Inc and is a paid consultant for Relay Therapeutics, Profluent Bio, Vilya Therapeutics, and Monimoi Therapeutics. All other authors declare no competing interests.

### Summary of Updates

Final revisions for elife - mostly renaming of figures.

